# Evaluation and Optimization of Motion Correction in Spinal Cord fMRI Preprocessing

**DOI:** 10.1101/2020.05.20.103986

**Authors:** Hamed Dehghani, Kenneth A Weber, Seyed Amir Hossein Batouli, Mohammad Ali Oghabian, Ali Khatibi

## Abstract

Motion correction is an essential step in the preprocessing of functional magnetic resonance imaging (fMRI) data, improving the temporal signal to noise ratio (tSNR) and removing unwanted variance. Because of the characteristics of the spinal cord (non-rigidity, surrounded by moving organs), motion correction becomes especially challenging. We compared the efficiency of different motion correction protocols and suggest a preferred method for spinal cord fMRI data. Here we acquired gradient-echo echo-planar-imaging axial lumbar spinal cord fMRI data during painful mechanical stimulation of the left lower extremity of 15 healthy volunteers on a 3T scanner. We compared multiple motion correction techniques: 2D and 3D FLIRT realignment with and without slice-wise regulation, SliceCorr (implemented in the Spinal Cord Toolbox) and proposed a method 3D FLIRT in addition to Slice Regulation (SLiceReg) along the spinal cord. TSNR, image entropy, DVARS, image Sum of Absolute Differences and number of activated voxels in the spinal cord from GLM analysis to evaluate the performance of multiple motion correction procedures. The tSNR and DVARS 3D FLIRT + SLiceReg were significantly improved over other realignment methods (p<0.001). In comparison, tSNR=14.20±0.02 and DVARS=165.77±1.54 were higher than other methods. Additionally, the number of activated voxels of the statistical map in our suggested method was higher than the other realignment methods (p<0.05). Our results illustrated the proposed motion correction algorithm that integrated 3D motion correction and 2D slicewise regularization along spinal cord curvature could improve subject-level processing outputs by reducing the effects of motions. Our proposed protocols can improve subject-level analysis, especially in lumbar region that suffers from involuntary motions and signal loss due to susceptibility effect more than other spinal cord regions.

## 1. Introduction

Motion correction is an essential processing step in fMRI in which the volumes of an fMRI time series are aligned to reduce the effect of participant movement during scanning and improve the sensitivity and specificity to detect blood oxygenation level-dependent (BOLD) signal changes (Oakes et al., 2005). For brain fMRI, motion correction approaches commonly employ six parameter rigid-body transformations (x, y, and z translations and rotations) to spatially realign each volume of the time series to a selected reference volume (e.g., the first, middle, or mean volume)(Bannister, Brady, & Jenkinson, 2004; Jenkinson, 2006; Kim, Boes, Bland, Chenevert, & Meyer, 1999). Rigid-body motion correction algorithms for the brain are generally sufficient as the shape of the brain remains largely constant over time with only the position and orientation changing due to movement.

Recent advancements in spinal cord image acquisition (e.g., reduced field-of-view imaging, advanced shimming procedures, and simultaneous brain-spinal cord acquisitions) and image analysis techniques have increased the utility and applications of spinal cord fMRI in the non-invasive study of spinal cord processing. The articulated structure of the spine and the surrounding anatomy present unique challenges for motion correction. While rigid body motion correction algorithms are largely accepted for brain fMRI, the non-rigidity of the spine leads to non-rigid deformations due to bulk motion, swallowing, and respiratory-induced motion; cardiac and CSF pulsations lead to non-uniform fluctuations in signal intensity across the image; and the respiratory cycle causes susceptibility changes in the lungs which induce B_0_ field distortions and shifts along the phase-encoding direction (Bannister et al., 2004; Julien Cohen-Adad et al., 2010; Leitch, Figley, & Stroman, 2010).

Previous spinal cord fMRI studies have employed a wide range of motion correction algorithms from conventional six parameter rigid-body motion correction algorithms to more sophisticated approaches to attempt to correct for the non-rigid motions. While motion correction in the spinal cord fMRI is ultimately important, it has yet to be studied systematically. The purpose of this study was to test the performance of different motion correction techniques using gradient-echo echo-planar-imaging (GE-EPI) axial lumbar spinal cord fMRI data during painful mechanical stimulation of the left lower extremity. We aimed to compare multiple motion correction techniques and suggest an optimal motion correction protocol.

## 2. Method

### 2.1. Subjects

Fifteen healthy volunteers (male, mean age ± standard deviation =25.88±4.44 years) participated in the study as a part of experiments to evaluate the feasibility of fMRI in the lumbar spinal cord. All of the participants provided informed consent. The study received approval from the human research ethics review board of Tehran University of Medical Sciences.

### 2.2. Data acquisition

All of the experiments were carried out using a 3 T whole-body MRI system (Siemens Magnetom Prisma; Siemens, Erlangen, Germany). Subjects were carefully positioned in the scanner with the longitudinal axis was parallel with the spinal cord. A three-plane localizer image was then taken to provide a survey reference. Radio-frequency (RF) pulses were supplied with a body coil, and a spine phased-array coil was used for receiving the signal in the lower thoracic-lumbar spinal cord.

Following the acquisition of the localizer reference image, the imaging was performed using the ZOOMit selective field-of-view (FOV) GE-EPI sequence with the following imaging parameters: TR/TE = 3000ms/30ms, FOV = 128×128 mm, matrix size = 128×128, in-plane resolution = 1×1 mm^2^, slice thickness = 3mm, and flip angle = 80 (Weber II, Chen, Wang, Kahnt, & Parrish, 2016). Spectral attenuated inversion recovery (SPAIR) was applied to suppress the effect of fat in GRE-EPI images. MRI images were acquired in the axial orientation and sampled in ascending interleaved order. The FOV spanned from the 9^th^ thoracic (T9) to the 2^nd^ lumbar (L2) vertebrae.

While scanning, painful mechanical stimuli were applied to the left foot (L5 dermatome) between two malleoli using a monotonic pressure device. For each participant, the subjective threshold for pain was calculated using the ascending staircase method (group mean = 3.9 ± 0.48 kg)(Gracely, Lota, Walter, & Dubner, 1988). Each block with a duration of 60s contained ten 3s duration painful stimuli 3s interval) at the participant’s threshold in 6 separated blocks which lasted a total of 540 seconds.

### 2.3. Image realignment methods

In spinal cord fMRI, especially in the lumbar spine, it should be considered that maximum movement was in z orientation (parallel to the superior-inferior axis of the spine) due to the diaphragm and renal movements. Some motions in the body can affect the spine the same as bulk motion like the movement of the larynx in the neck, displacement of internal organs in the chest and abdomen following the diaphragm movement. Either direction of these non-rigid motions and orientation of the spine is a crucial parameter to determine optimum realignment technique. We applied some of the previously recommended methods and propose a novel technique based on previous work by our team. It should noted that some of these methods limit the motion correction to a specific region in the image. A binary mask based on spinal cord segmentation was automatically generated (sct_create_mask) and used for the purpose of motion correction. The centerline of the spinal cord mask was then used to create a cylindrical mask with a radius/diameter of 45 voxels. This mask was used during motion correction to weight the reference image and exclude areas outside of the vertebral column.

#### 2.3.1. Default 3D Volume Correction with MCFLIRT

MCFLIRT is a rigid-body motion correction tool based on affine registration tool in FSL toolbox. This tool by default uses the middle volume as a reference image and identifies/calculates the transformation between the reference image and each volume in the time series. This method uses normalized correlation as a *default* cost function, six DOF for slice transformation process and b-spline interpolation to obtain robust rigid registration (Jenkinson, 2006; Smith et al., 2004).

#### 2.3.2. Default motion correction with 2D Volume Correction with FLIRT

Due to the specific structure of the spinal cord, some researchers have proposed that since the z-axis movement is very limited, a 3D motion correction seems unnecessary. Yet, in a recent paper, motion correction along the z-axis with 2D and 3D co-registration is recommended for the first step of motion correction (Keliris et al., 2007; Weber II et al., 2016). We included this method to test this assumption and compare the outcome with the 3D correction methods.

#### 2.3.3. Slicecorr

This method is based on 2D rigid-body realignment that assumes the spinal cord has no rostrocaudal bulk motion (i.e. along the z-axis) and no rotation around x and y directions. Slicecorr uses correlation as the cost function, three DOF transformation per slice, fmeansearch as Minimization algorithm and mean volume image as a reference template (J Cohen-Adad, Rossignol, & Hoge, 2009). Slicecorr is the default motion correction protocol for the spinal cord toolbox (SCT). SCT fMRI motion correction uses a previously created mask to decrease the effect of organs that move independently from the spine. This binary mask is used in the motion correction algorithm to weight the cost function metric to the spinal cord (De Leener et al., 2017).

#### 2.3.4. 3D Volume Correction + 2D Slice-wise Correction

In the first step of this motion correction method, a 3D rigid-body realignment using normalized correlation as a cost function and b-spline interpolation has performed. A cylindrical binary mask, created from spinal cord centerline with automatic segmentation of cord on the mean volume image by SCT, can improve 3D rigid-body realignment on the spinal cord. Then the mean of the previous output is applied as the reference image in the second step. In this step, the 3D motion corrected time series is entered to perform 2D slice-wise realignment for each slice and remove the effect of non-rigid motions (Summers, Brooks, & Cohen-Adad, 2014; Weber II et al., 2016).

#### 2.3.5. 3D motion correction (FLIRT)+ 2D Slice-wise realignment and regularization (SliceReg)

Referring to the previous method, we assumed that motion correction with two steps could be improved by updating registering metrics for improving the efficacy of realignment procedure and maintaining the structure of functional images. We assessed the multiparameter affine rigid-body registration methods in the spinal cord images and checked the signal to noise ratio (SNR) for them. Based on this experiment we suggested in the first step, least-square as an efficient cost function, b-spline as interpolation metric and use of the mean volume of non-motion corrected time series as the reference image (Barakat et al., 2012; Middleton et al., 2014). Again, we used a binary mask to cover the spinal cord and exclude other organs in the axial slices. In the second step, the output of 3D-realignment was entered in the 2D slice-wise realignment procedure. Mean squares was used as the cost function for the curvature of lumbar spine and spline as interpolation can lead to the best result in the registration function. SliceReg estimates slice-by-slice translations and regularization qualifications in the Z-axis direction that can improve the second step of motion correction (J Cohen-Adad, Levy, & Avants, 2015; De Leener et al., 2017). The preferred motion correction step was performed by freely available toolboxes (FSL and SCT) as summarized in Table 1.

**Table 1.**
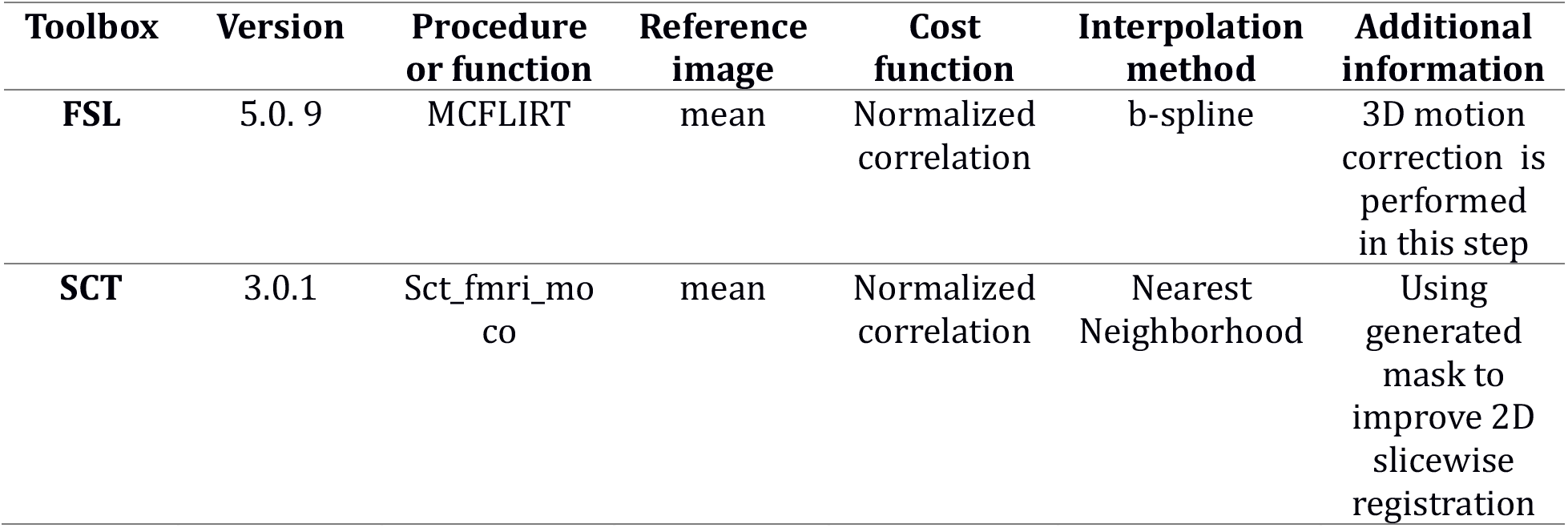
Motion correction parameters and software packages

| **Table 1**<br>**Motion correction parameters and software packages** |  |  |  |  |  |  |
| --- | --- | --- | --- | --- | --- | --- |
| **Toolbox** | **Version** | **Procedure or function** | **Reference image** | **Cost function** | **Interpolation method** | **Additional information** |
| **FSL** | 5.0.9 | MCFLIRT | mean | Normalized correlation | b-spline | 3D motion correction is performed in this step |
| **SCT** | 3.0.1 | Sct_fmri_moco | mean | Normalized correlation | Nearest Neighborhood | Using generated mask to improve 2D slicewise registration |

### 2.4. Image Analysis Motion Correction Performance

To evaluate the effectiveness of motion correction methods and their registration parameters, several performance metrics were calculated and compared.:

#### 2.4.1. Temporal Signal to Noise Ratio (tSNR)

In addition to MRI background noise, addiitonal components of noise in fMRI include noise arising from the subject (i.e. cardiac and respiratory pulsations, movements) and task related noise (Tong, Hocke, & Frederick, 2019; Triantafyllou, Polimeni, & Wald, 2011). In fMRI, fluctuations in the BOLD signal are measured over time. Motion correction algorithms reduce noise due to bulk motion across the time series. Temporal SNR (tSNR is used to characterize the stability of the BOLD signal over the fMRI time series And was used as our primary outcome measure of motion correction performance. Changes in tSNR were assessed only in the spinal cord using a manually drawn binary spinal cord mask from the mean motion-corrected image. We calculated tSNR as the mean signal over time divided by the standard deviation of images over time using the following equation:

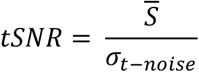

#### 2.4.2. Information Entropy

Information entropy is a measure of the amount of randomness (or uncertainty) there is in a signal or image; or more precisely, how much information is produced by the signal. Application of this measure without using a predetermined region of interest in the image has been suggested to be useful for optimizing the entropy of an image with low resolution in the phase encode direction which can provide a sufficiently accurate measure of motion (Atkinson et al., 1999). Entropy was calculated using the following equation:

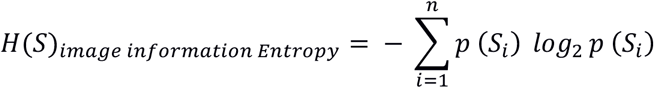

#### 2.4.3. DVARS

DVARS (D, temporal Derivative of time courses, VARS, variance over voxels) demonstrates the rate of signal change across the spinal cord at each frame of data. In an ideal data series, a value of DVARS depends on the temporal standard deviation and the temporal autocorrelation of the data (Nichols, 2017). To calculate DVARS, changes in each voxel’s value at each time point is compared to the previous time-point (Power, Barnes, Snyder, Schlaggar, & Petersen, 2012). DVARS was calculated in the whole image to find a metric that demonstrated standard deviation of temporal difference images in the 4D raw-data (Nichols, 2017). The value of DVARS, show how much the intensity of voxels within the spinal cord in each slice and at each volume change in comparison to the previous volume (Power et al., 2014). DVARS was calculated using the following equation:

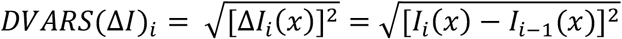

In this equation, Δ*I_i_*(*x*)is used as local image intensity on the frame. DVARS could result in more accurate modelling of the temporal correlation and standardization because it is obtained by the most short-scale changes (Nichols, 2017).

#### 2.4.4. Sum of Absolute Differences (SAD)

Sum of Absolute Differences (SAD) is a metric of the similarity of images in a series that is used as a motion estimator in image processing algorithms including MRI and fMRI quantitative analyze (Seshamani et al., 2013; Shakil, Keilholz, & Lee, 2016; Turk et al., 2017). To calculate this metric for estimating total motion, the absolute difference is considered between each voxel in the image series and the corresponding voxel in the next (Niitsuma & Maruyama, 2010; Vassiliadis, Hakkennes, Wong, & Pechanek, 1998).

### 2.5. fMRI Data Analysis

#### 2.5.1 Physiological Noise Correction

In the fMRI studies, physiological noise is described as cardiac and respiratory-related changes in the signal which can influence the analysis of fMRI data. This includes the motion of the spinal cord due to cerebrospinal fluid (CSF) pulsation and magnetic field fluctuations due to movement of lungs and changes in the air volume in the thorax and lungs (Dehghani et al., 2020). In order to minimize the effect of physiological noise, we followed a two-step correction protocol:

##### 2.5.1.1. ICA-based correction

Spatial Independent Component Analysis (ICA) can classify signal components as neural-related or noise-related and remove noise to increase the statistical power of the analysis of fMRI data. This step was performed in two levels, visual and quantitative classification of independent components. In the visual classification, the location of activated clusters (and their peaks) in the spinal cord suggests the neural-related origin of the component, whereas clusters mainly located in the vertebra, discs, blood vessels (main arteries) and especially in the kidneys are usually correlated to physiological noise (respiration, pulsation). Sometimes the structure of non-physiological patterns related to the MRI sequence, hardware artefacts, or interactions of the acquisition with physiological motion is seen in the images of some component (Griffanti et al., 2017).

In the quantitative classification, components were obtained for each data set, which had the significance of the BOLD signal variability. The components with specified criteria were determined as noise: (a) the power of the spatial component’s time series at high frequencies were larger than 0.08 Hz (b) more than 50% of significantly activated voxels [Z > 2.3] was seen out of the spinal cord mask in the component’s spatial map. Overall, 5 to 16 components have both considered criteria in subjects, and their time series and components were filtered as a noise before (Kelly Jr et al., 2010; Vahdat et al., 2015).

##### 2.5.1.2. Physiological Noise Modelling

Respiratory signals, cardiac signals, and MRI triggers are acquired using Siemens Prisma scanner sensors (pulse oximeter sensors and respiratory cushion) and Physiological Monitoring Unit (PMU) (sampling rate = 400 Hz). To remove residual effects of physiological noise, slice-specific noise regressors were generated using a custom-made physiological noise modelling tool which was adapted to PMU recording signals. This MATLAB custom-made tool uses a model-based approach similar to the Retrospective Image Correction (RETROICOR) and FSL’s Physiological Noise Modelling tool (PNM) as described by Glover et al. and Brooks et al.(Brooks et al., 2008; Glover, Li, & Ress, 2000). After down-sampling of the physiological signal, a cardiac phase and respiratory phase and the first three harmonics were assigned to each volume and regressors were created which can be used to model the physiological noise within the General Linear Model (GLM).

#### 2.5.3. General Linear Model analysis for preprocessed fMRI data-sets

First, the spinal cord was extracted from the fMRI images using an automatic drawn spinal cord mask that was generated via Spinal Cord Toolbox (SCT). Motion-corrected images from the output of different motion correction methods were concatenated into an fMRI time series. Then slice-timing correction and image intensity normalization was performed. After that advanced spatial smoothing was performed with a Gaussian kernel of 2 mm full width half maximum (FWHM) in the straight spinal (sct_straighten_spinalcord) cord and high pass temporal filtering (sigma = 90 s). FSL’s motion outlier detection tool was then applied to detect outlier time points using DVARS and box-plot cutoff = 75th percentile + 1.5 × interquartile range (IQR) for thresholding (Power et al., 2012; Weber II et al., 2016).

FMRIB’s Improved Linear Model (FILM) with prewhitening were used to generate statistical maps of the preprocessed data for each subject (Woolrich, Ripley, Brady, & Smith, 2001; Worsley et al., 2002). The design matrix contained the standard hemodynamic response function convolved pressure pain stimulation vectors (included HRF parameters from FSL), the physiological noise vectors, and the temporal masks of outlier time points were included as covariates of no interest. Voxels with a p < 0.05 (uncorrected) were considered active in the spinal cord mask.

In order to eliminate the effect of normalization steps and the linear and non-linear registration in the higher-level analysis, we decided to perform GLM analysis just at thesubject level. Investigating the effects of motion correction at the subject level allows us to focus on the the effects of the motion correction methods more specifically without introducing other potential sources of variance.

### 2.6. Statistical Analysis

The motion correction performance metrics (tSNR, Entropy, DVARS, Sum of Absolute Differences and the number of activated voxels) were calculated and compared across the different motion correction methods. The mean of each parameter was computed. Statistical analyses were performed using SPSS (Version 16.0, The SPSS, Inc., Chicago, IL, USA) and CRAN (R package, Version 3.5.1, R Development Core Team). Normality of the metrics was assessed with the Kolmogorov-Smirnov test. The mean of normal data (uncorrected as well) for each method was processed with One-way ANOVA with repeated measures in within-subjects comparison and then multiple comparison post-hoc test with Bonferroni correction was performed for statistically significant results Partial eta squared (n_p_^2^ is reported to present the estimation of the effect size for significant results). Values of 0.01, 0.06 and 0.15 represent small, moderate and large effect sizes respectively. To assess the relationship among parameters, Pearson Correlation test was applied in the optimized motion correction algorithm, which can both explain the effect of motion correction on parameters and that of each parameter on the quality of the image. Finally, utilizing the results of this test, we will be able to choose the best parameter for motion outlier volumes detection and scrubbing it in the subject level GLM data processing.

## 3. Results

As expected, generally, motion correction improved the quality of the data as indicated by improvement in each of the parameters. In table 3, the results of descriptive statistical data analysis are presented. A one-way repeated measures ANOVA was applied to compare the effect of motion correction algorithms on the uncorrected (raw data) in 2D and 3D FLIRT motion correction, SCT fMRI motion correction, FLIRT 3D and 2D+ silcewise and FLIRT 3D + SliceReg motion correction method. Figure 1 presents an overview of different measures on different motion correction methods.

**Table 2.** Summary of Image Analysis parameters between different motion correction methods (df=6)

| Subset for alpha=.05 |  |  |  |  |
| --- | --- | --- | --- | --- |
| parameters |  | Mean | F value | p value |
| tSNR | Uncorrected | 9.343 | 63.2 | <0.001 |
|  | FLIRT3D | 13.871 |  |  |
|  | FLIRT2D | 12.854 |  |  |
|  | FLIRT2D+Slicewise | 12.873 |  |  |
|  | FLIRT 3D+Slicewise | 13.432 |  |  |
|  | SCT motion correction | 9.154 |  |  |
|  | FLIRT 3D+SliceReg | 14.200 |  |  |
| Entropy | Uncorrected | 0.721 | 0.998 | 0.441 |
|  | FLIRT3D | 0.701 |  |  |
|  | FLIRT2D | 0.698 |  |  |
|  | FLIRT2D+Slicewise | 0.708 |  |  |
|  | FLIRT 3D+Slicewise | 0.708 |  |  |
|  | SCT motion correction | 0.729 |  |  |
|  | FLIRT 3D+SliceReg | 0.702 |  |  |
| **DVARs** | Uncorrected | 182.640 | 4.13 | 0.001 |
|  | FLIRT3D | 150.453 |  |  |
|  | FLIRT2D | 155.234 |  |  |
|  | FLIRT2D+Slicewise | 191.163 |  |  |
|  | FLIRT 3D+Slicewise | 165.772 |  |  |
|  | SCT motion correction | 190.927 |  |  |
|  | FLIRT 3D+SliceReg | 165.775 |  |  |
| **SAD** | FLIRT3D | 1.343 | 22.95 | <0.001 |
|  | FLIRT2D | 1.919 |  |  |
|  | FLIRT2D+Slicewise | 1.542 |  |  |
|  | FLIRT 3D+Slicewise | 0.648 |  |  |
|  | SCT motion correction | 0.829 |  |  |
|  | FLIRT 3D+SliceReg | 0.501 |  |  |
| **Activated voxels** | Uncorrected | 28.933 | 22.95 | 0.002 |
|  | FLIRT3D | 38.266 |  |  |
|  | FLIRT2D | 34 |  |  |
|  | FLIRT2D+Slicewise | 48.2 |  |  |
|  | FLIRT 3D+Slicewise | 52.466 |  |  |
|  | SCT motion correction | 29.333 |  |  |
|  | FLIRT 3D+SliceReg | 58.333 |  |  |

**Table 3.** Results of Pearson correlation between outcome measures on the optimized motion correction method

|  | 1 | 2 | 3 | 5 |
| --- | --- | --- | --- | --- |
| 1. Difference Intensities |  |  |  |  |
| 2. tSNR | 0.36 |  |  |  |
| 3. Entropy | -0.27 | -0.51 |  |  |
| 5. DVARS | -0.10 | -0.30* | 0.03 |  |
| * Demonstrate significant $r$ coefficient $p<0.05$ | | | | |

In the comparison of temporal SNR, the average tSNR ± standard error (SE) significantly increased from 9.41 ± 1.51 arbitrary units (AU) (figure 2). There was a statistically significant effect of motion correction methods on tSNR parameter, *F*(6,84) = 63.2, *p*<.001, 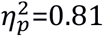.

**Figure 2.**
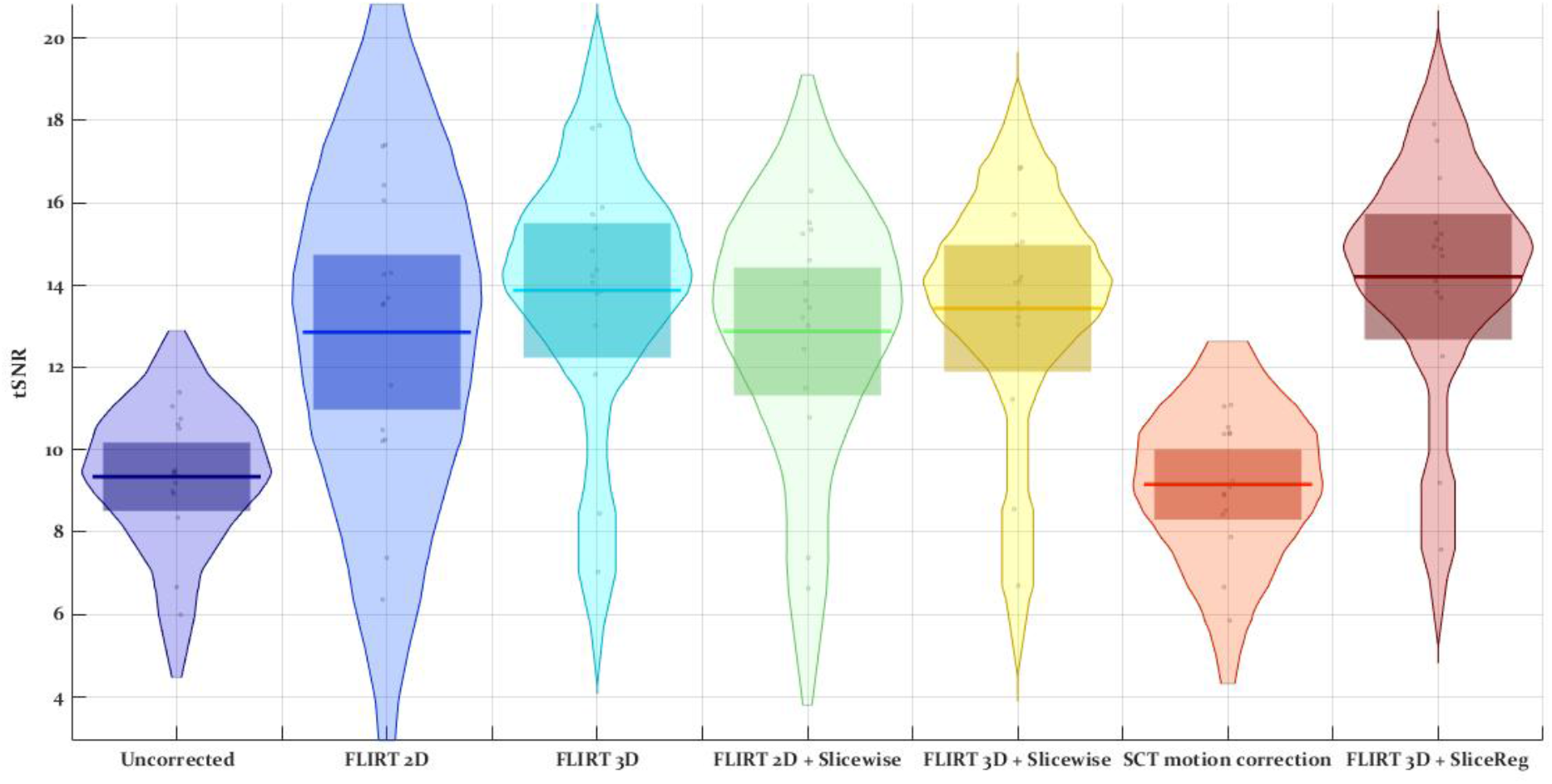
Summary of temporal SNR of motion-corrected images from proposed motion correction methods

FLIRT 3D + Slicewise had the maximum single voxel tSNR= 26.41. Post hoc multiple comparisons using the Bonferroni correction indicated that the mean tSNR for the FLIRT 3D + Slicereg was significantly greater than the other motion correction methods and raw-data (*p*<.01) except for the FLIRT 3D motion correction algorithm (*p*=.578).

*In the comparison of the entropy of information measure, a statistically significant effect could not be seen* (*F*(6,84) =0.998, *p*=.441). None of motion correction methods had a significant effect on the entropy of uncorrected image.(0.721±0.005) (figure 3).

**Figure 3.**
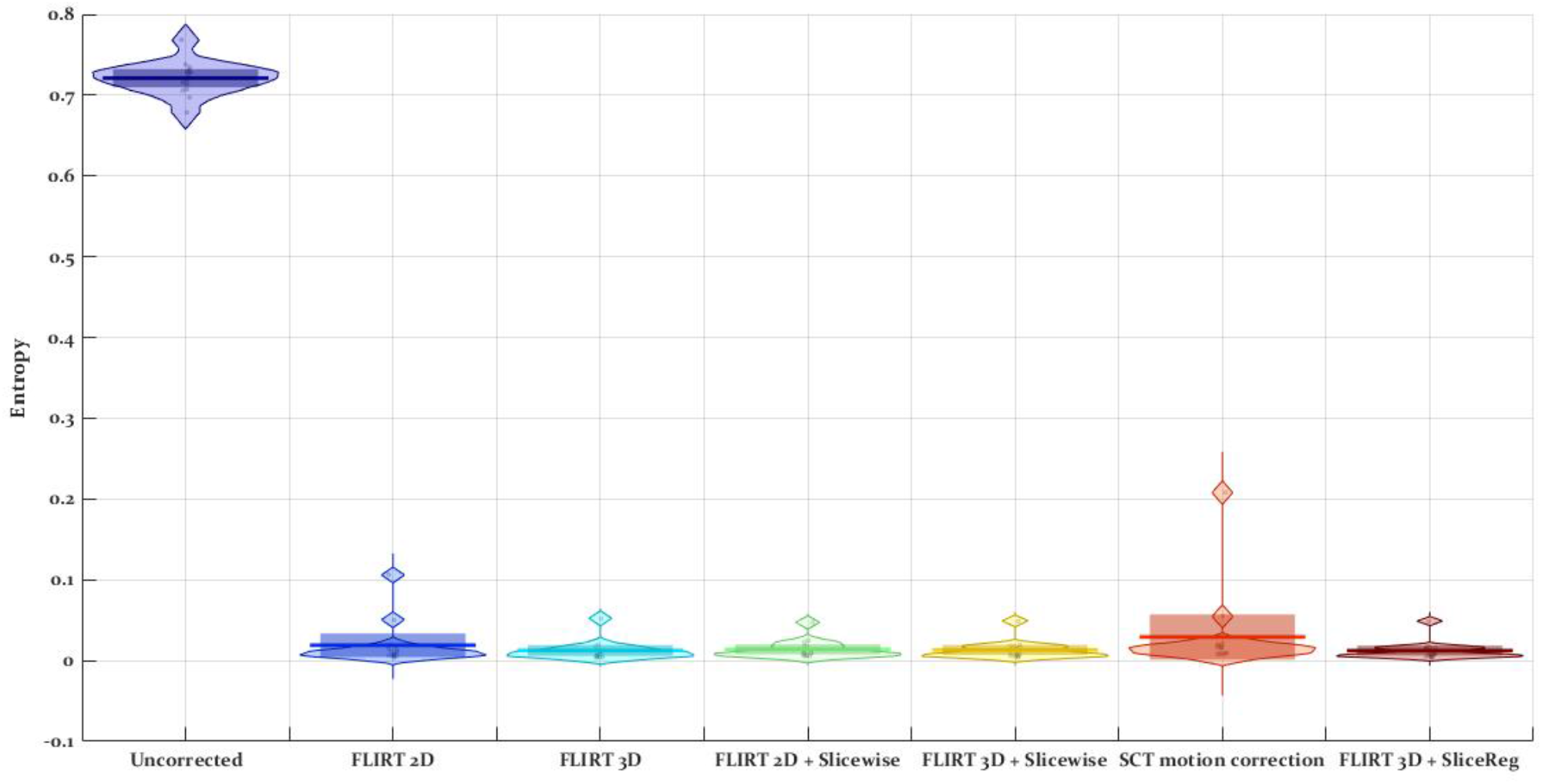
Summary of temporal Entropy of motion corrected images from Proposed motion correction methods.

**Figure 4.**
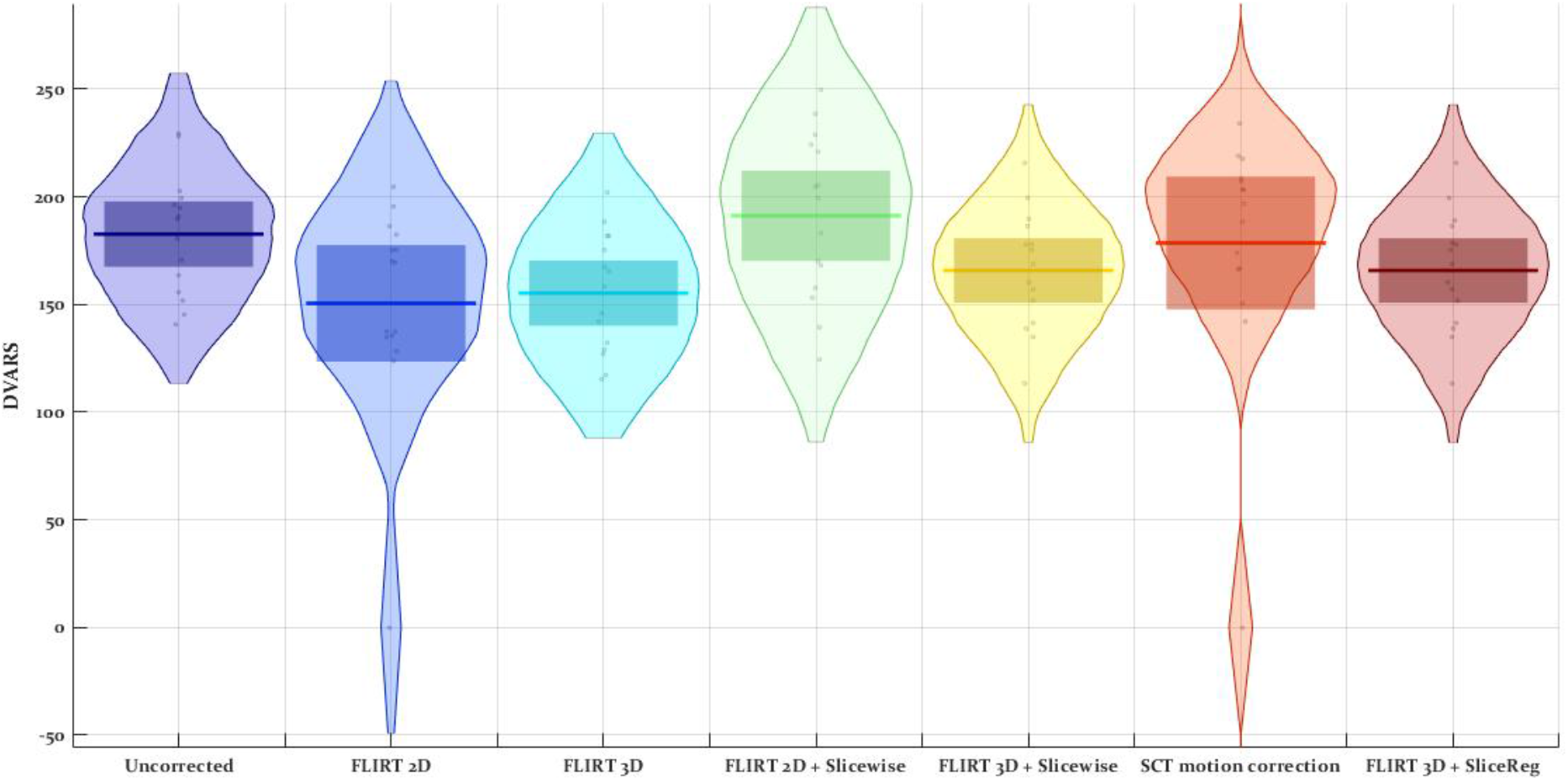
Summary of DVARS parameter in motion corrected images from Proposed motion correction methods

*There was a statistically significant effect of motion correction methods on DVARS parameter, F*(6,84) = 4.01, *p*<.001, 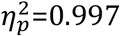. DVARS parameter demonstrate RMS intensity difference of volume by volume effect of the motion correction 3D FLIRT method has minimum error 150.45 ± 28.43 and this value in the proposed motion correction method is 165.77 ± 1.54 (Figure 5). Post hoc multiple comparisons using the Bonferroni test indicated that the mean of DVARS metric for the FLIRT 3D + Slicereg was significantly different than the uncorrected raw-data, Flirt 2D+ slicewise and SCT motion correction algorithm (*p*<.05).

**Figure 5.**
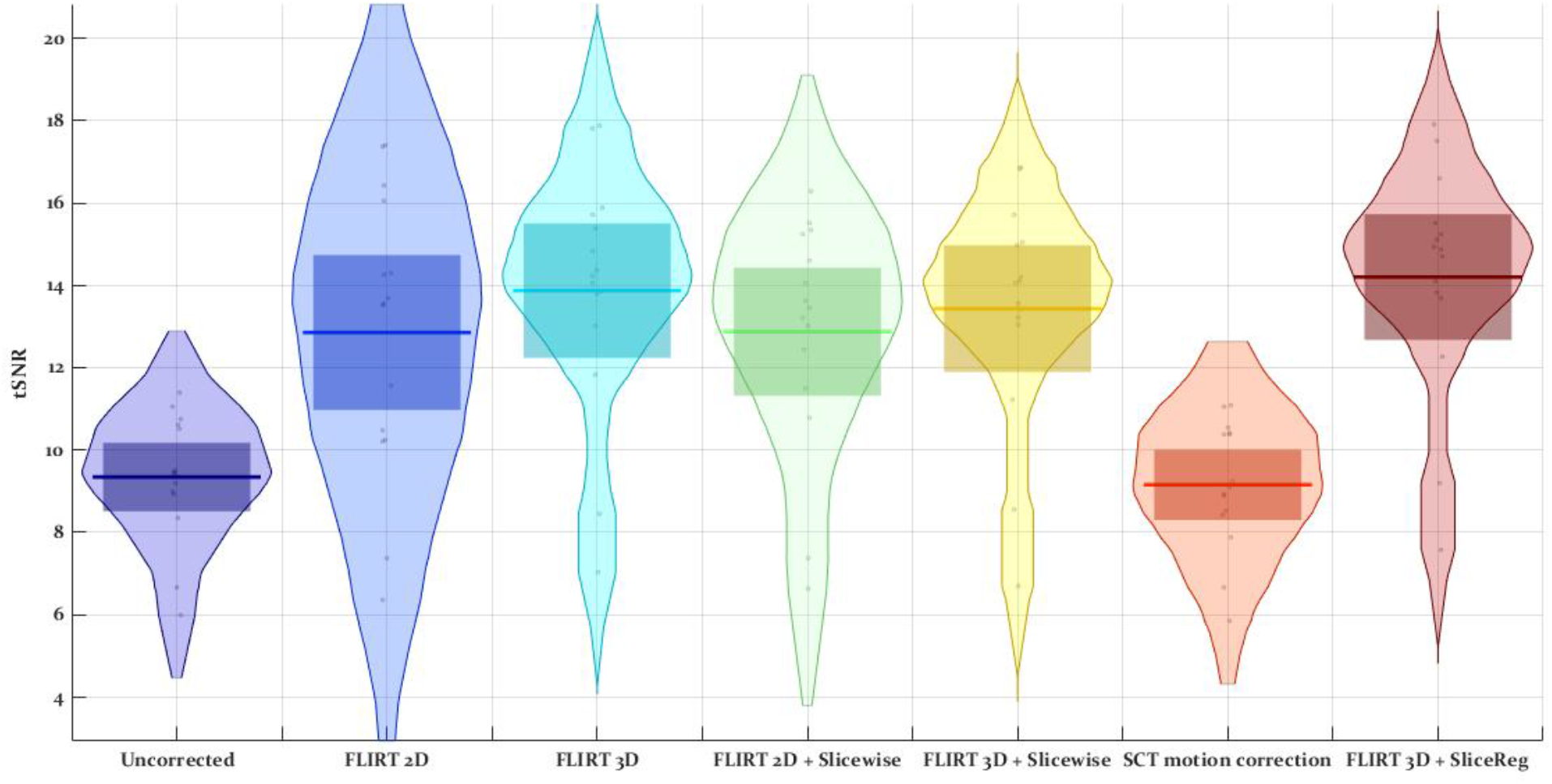
Summary of Activated voxels (in the Spinal cord) in motion corrected images from Proposed motion correction methods

After subject level GLM, the number of activated voxels of the statistical map was summed in the spinal cord, then compared using repeated measures ANOVA (Figure 7 and 8). There was a significant effect of motion correction method on the sum of the activated voxels, F(6, 84) = 22.95, p=0.002, 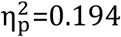. The number of activated voxels without any motion correction was 28.93±24.89 and with 3D+2D silcewise motion correction was 58.33±40.21.

**Figure 8.**
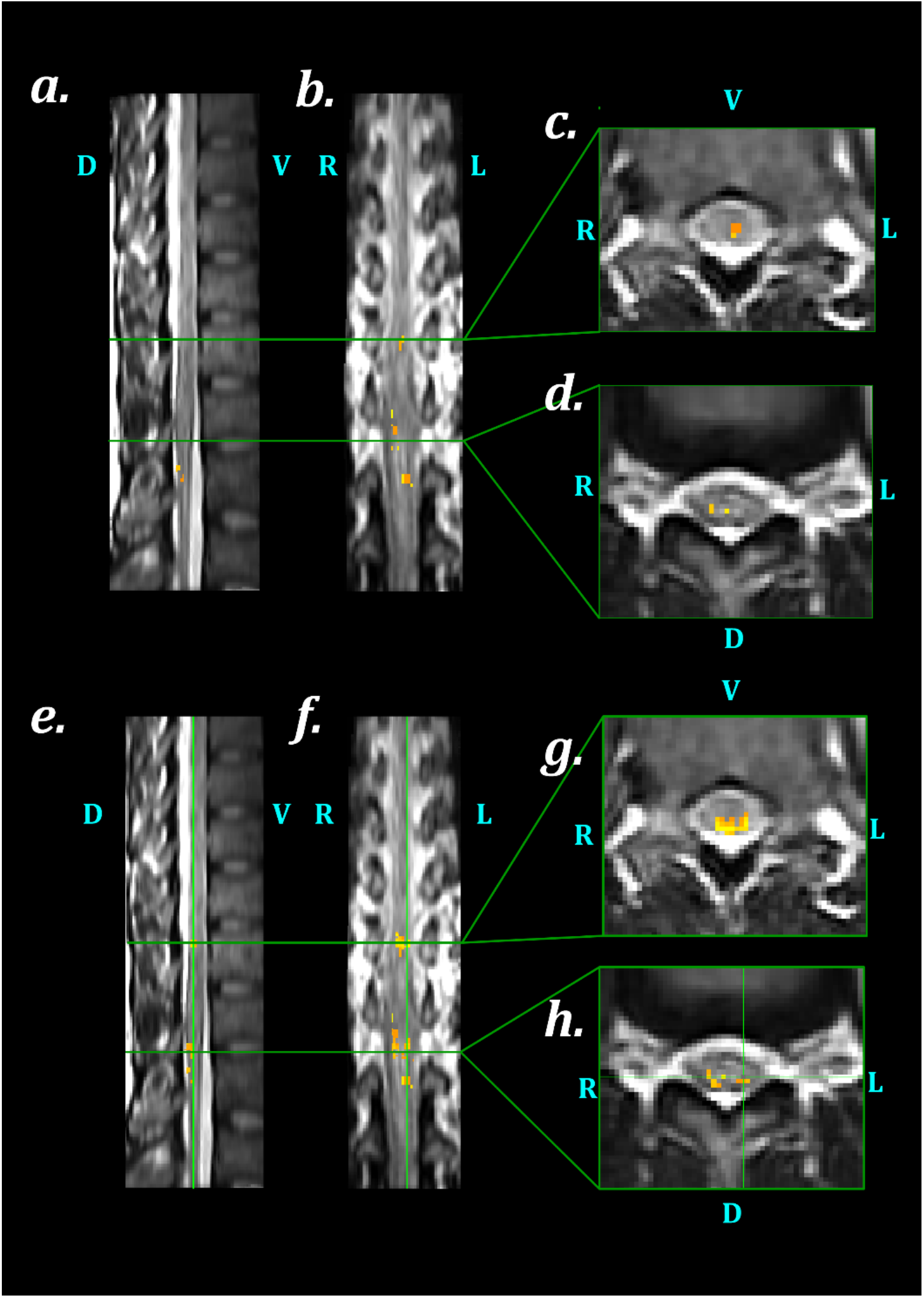
Comparison of Activation map of spinal cord fMRI GLM analysis based on induced pressure pain (3.9 ± 0.48 kg), by considering two Motion correction methods. (a), (b), (c), (d) illustrate activated voxels in the spinal cord on three views (Sagittal, Coronal and Axial), based on 3D FLIRT Motion correction; (e), (f), (g) and (h) illustrate activated voxels in the spinal cord on three views (Sagittal, Coronal and Axial), based on the proposed Motion correction (3D FLIRT slice-wise + SliceReg 2D slice-wise).

**Figure 9.**
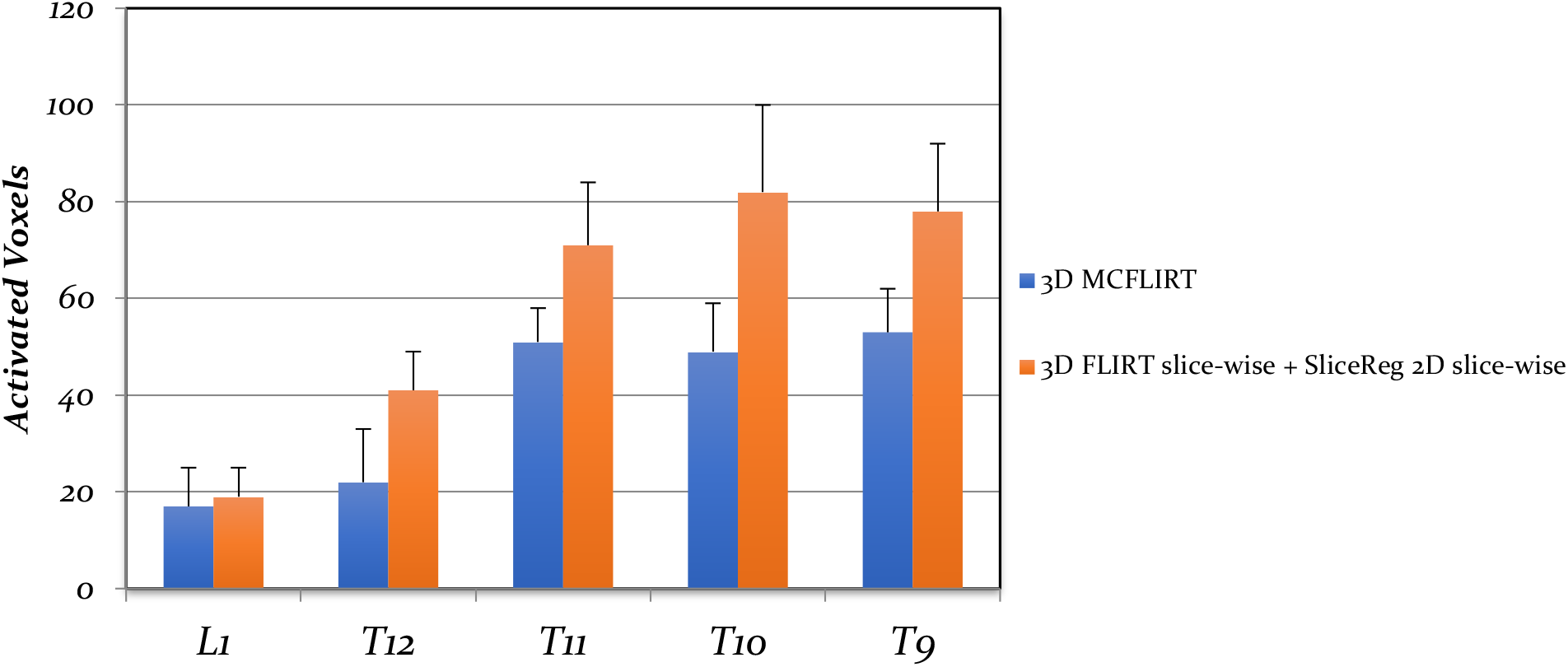
Within subjects comparison of Activated voxels of fMRI GLM analysis in spinal cord levels, by considering two Motion correction methods. A comparison between Activated voxels of two GLM analysis based on two motion correction methods (3D FLIRT and 3D FLIRT slice-wise + SliceReg 2D slice-wise) illustrates optimized motion correction methods increase the activated voxels in the Thoracic vertebral level of the spinal cord, nearby moving diaphragm and lungs.

Post hoc multiple comparisons using the Bonferroni correction indicated that the mean of Activated voxels for the FLIRT 3D + Slicereg was significantly different than the uncorrected raw-data, FLIRT 2D, FLIRT 3D and SCT motion correction algorithm (*p*<.05).

A Pearson product-moment correlation coefficient was computed to assess the relationship between the image analysis metrics in the optimized motion correction method in order to come up with suggestions for DVARS as a metric to censor motion outlier time points. The result of Pearson Correlation test was summarized in Table 4. It is clear that there was a negative correlation coefficient between DVARS and tSNR, r =-0.303, n = 15, *p*<0.05. This significant correlation coefficient illustrates with increase tSNR parameter and decreases DVARS.

## 4. Discussion

In this study, the effectiveness of different motion correction algorithms was tested using GRE-EPI axial lumbar spinal cord fMRI datasets during painful pressure stimulation of the left lower extremity. Several reported motion correction algorithms were studied, algorithms using a combination of rigid and non-rigid body registrations were more effective in motion correction leading to greater improvemetns in the image quality parameters. The effect of the motion correction was then explored on the activity in the cord, and again, the algorithms using a combination of rigid and non-rigid body registrations resulted in greater cord activation. Of the motion correction methods tested, the 3D FLIRT motion correction andslice-wise algorithm consistently outperformed the other algorithms and is recommended for use in future studies.

In comparison with preprocessing steps for the fMRI data acquired from the brain, no specific pipeline exists for data preparation in the statistical processing in the spinal cord fMRI. A major step in this process is motion correction which has been suggested to be very important in determining the output of fMRI studies. Existing studies either used in-home developed algorithms adapted for the spinal cord data (Lawrence, Kornelsen, & Stroman, 2011; Rempe et al., 2015), used tools not specifically designed for the spinal cord, or used SCT. Surprisingly and despite the growing number of spinal cord fMRI studies, no study to date has systematically investigated the effect of the different motion correction algorithms on the quality of the spinal cord fMRI signal and the level of spinal cord activation.

The optimized motion correction algorithm (FLIRT 3D+Slicerg, see section 2.3.5.) was performed in two steps; in the 1^st^ step, 3D motion correction was applied with FLIRT registeration algorithm and binary mask to exclude out of vertebrae tissues; and in the 2^nd^ step the output of previous step was entered in 2D slice-by-slice realignment procedure and regularization along the spinal cord using SliceReg. The optimized motion correction algorithm could minimize the effect of voluntary bulk motion and non-voluntary physiological movements. It increased tSNR value with almost a rate of 1.5 times more than other motion correction methodologies considering the main image (uncorrected) tSNR. Quantitative measures such as DVARS used to evaluate the effect of spinal cord movement and the correction protocols on the results of GLM subject-level analysis. Interestingly, our results suggest that FLIRT 2D is slightly better in performance regarding this measure. Sum of absolute differences, used as an motion estimator and similarity measure in the registration algorithm, illustrates the correlation between tSNR and other related parameters. The highest correlated parameters are tSNR and RMSE with R=-0.784. These parameters are utilized as quality control measures in fMRI raw-data (Marcus et al., 2013)-

Information entropy can be defined as how much randomness (or uncertainty) there is in a signal or an image; information entropy provides a measure of how much information is provided by the signal or image. The result of ANOVA test demonstrate information entropy differences in motion correction methods was not statistically significant. Based on information entropy theory concept, Subject and medical scanner conditions make the most impact on entropy of images and image processing steps has the least impact (Tsai, Lee, & Matsuyama, 2008).

Although Frame-wise displacement, which is used to regress out motion outliers time points, does not consider rotation movements leading to underestimation of this method, therefore calculating DVARS parameter is the best way to detect and scrub changes in motion outlier signal (Caballero-Gaudes & Reynolds, 2017; Ciric et al., 2017). DVARS measure the changes in image intensities comparing to the previous time point in a time sires voxel-wise (or ROI-wise) as opposed the global signal which is the average intensity of at a time (Power et al., 2014; Satterthwaite et al., 2013), and the average value of DVARS depends on the temporal standard deviation considered in tSNR formula (Nichols, 2017). The negative correlation between DVARS and tSNR demonstrates the fact that any reduction in DVARS results in an increase in tSNR. The acceptable value of DVARS varies over localities, making it difficult to create comparable summaries of data qualities in data series (Afyouni & Nichols, 2018). Examination of the regional variation caused by motion, in the BOLD signal, provides some novel insights into the nature of motion-BOLD relevancy which appear in the reflection of motion-induced artefacts, while others are driven from motion-related changes in neural activity.

The optimized motion correction method presented here (including FLIRT 3D motion correction + SliceReg) may improve the ability of detection of activated voxels based on the accepted tSNR, DVARS in the preprocessing step of the analysis. Optimal tSNR, DVARS and RMSEs may lead to stronger trends of higher z-scores and consequently more accurate active voxels (Van Der Zwaag, Da Costa, Zürcher, Adams, & Hadjikhani, 2012).

### 4.1. Limitations and Future Work

Our results in this study depend critically on the quality of MR images. Quality of MR images relies on some parameters such as image resolution (matrix, field of view, slice thickness), region of interest in imaging procedure, can affect the spinal cord fMRI preprocessing steps like co-registration and segmentation (Sabaghian, Dehghani, Batouli, Khatibi, & Oghabian, 2020). The field of view and orientation of raw-data was axial and all of preprocessing steps are dedicated to axially oriented raw-data. Physiological micromovements and effect of changing magnetic susceptibility during respiration also influence the motion correction and result of the fMRI single and group processing. Some previous studies performed spinal cord fMRI in the sagittal orientation with SE-HASTE sequence. It should be noted that the current method may not be optimal for sagittal or coronal acquisitions. Besides, the cut-offs for DVARS should be investigated in future studies for motion scrubbing.

### 4.2. Conclusion

This paper aimed to expand knowledge about spinal cord fMRI imaging and motion artifact removal in several methods. We propose a motion correction algorithm that integrated 2D slice-wise regularization and 3D motion correction can improve subject-level processing results by eliminating extended motions. A 3D motion correction with Mean Square cost function, 6 rigid-body degrees of freedom and linear interpolation (not systematically tested) is proposed for the first step of motion correction. A 2D slice-wise and regularization along spine curvature, Correlation as a cost function and linear interpolation algorithm are suggested for the second step. Creating appropriate mask based on the curvature of the spine can increase the accuracy of motion correction method and reduce the effect of out of spine movements. Future researches on spinal cord fMRI preprocessing method will contain improved instructions of artefact avoidance and removal.

## Acknowledgement

Authors gratefully acknowledge the use of the services and facilities of the National Brain Mapping Laboratory (NBML) in Tehran, Iran. We also would like to thank Shahabeddin Vahdat, Elaheh Sadri, Soodeh Moallemian, Elaheh Saleh for helpful discussions. This project was supported by the Tehran University of Medical Sciences Grant No. 94-03-30-29965.

